# Physiological traits and their relationships vary along an elevational gradient within and among Fijian bee species

**DOI:** 10.1101/2022.07.27.501487

**Authors:** Carmen RB da Silva, Julian E Beaman, Marika Tuiwawa, Mark I Stevens, Michael P Schwarz, Rosalyn Gloag, Vanessa Kellermann, Lesley A Alton

**Affiliations:** School of Natural Sciences, Macquarie University, North Ryde, New South Wales, Australia; School of Biological Sciences, Monash University, Clayton, Victoria, Australia; College of Science and Engineering, Flinders University, Bedford Park, South Australia, Australia; The University of the South Pacific, Herbarium, Viti Levu, Fiji; Earth & Biological Sciences, South Australian museum, Adelaide, South Australia, Australia; School of Biological Sciences, University of Adelaide, South Australia, Australia; School of Life and Environmental Sciences, University of Sydney, Sydney, New South Wales, Australia; School of Agriculture, Biomedicine and Environment, La Trobe University, Melbourne, Australia; Centre for Geometric Biology, Monash University, Clayton, Victoria, Australia

**Keywords:** metabolic cold adaptation, Krogh’s rule, discontinuous gas exchange, hygric hypothesis, bees, thermal performance, metabolic rate, frequency of gas exchange, functional traits

## Abstract

1. Temperature and water availability are hypothesised to be important abiotic drivers of the evolution of metabolic rates and gas exchange patterns, respectively. Specifically, the metabolic cold adaptation hypothesis (MCA) predicts that cold environments select for faster metabolic rates to counter the thermodynamics of biochemical reactions while the hygric hypothesis predicts that dry environments select for discontinuous gas exchange to reduce water loss.
2. Although these two hypotheses consider different physiological traits and how they vary along different abiotic gradients, metabolic rate drives frequency of gas exchange patterns in insects meaning these two traits are inherently linked. Despite this link, the MCA and hygric hypotheses are rarely considered together and the extent to which metabolic rates and frequency of gas exchange vary and co-vary across climatic gradients remains unclear.
3. We tested the MCA and hygric hypotheses within a species of endemic Fijian bee, *Homalictus fijiensis*, across an altitudinal gradient of 1100 m, and among four Fijian bee species, including *H. fijiensis*, that inhabit different altitudinal bands. In Fiji, environmental temperature is ∼5°C lower in the central highlands than in the coastal lowlands with the highlands receiving ∼100 mm of additional precipitation than the lowlands each month.
4. We found an MCA–like pattern within *H. fijiensis* and among Fijian bee species, where metabolic rate decreased with increasing temperature, but precipitation also explained variation in metabolic rate. However, we did not find support for the hygric hypothesis within *H. fijiensis* or among species (frequency of gas exchange was not negatively correlated with precipitation).
5. The relationship between metabolic rate and frequency of gas exchange was steeper for species that occupied lower elevations on average, suggesting it is possible that these two traits can evolve independently of each other despite being positively correlated.

## Introduction

Variation in climate shapes species’ physiological traits and geographic distributions (Diamond & Martin, 2021; Hoffmann & Sgro, 2011; Kellermann et al., 2012). For terrestrial ectotherms in particular, environmental temperature and aridity are hypothesised to be important abiotic drivers of the evolution of key physiological traits such as metabolic rate and respiratory patterns (Chown et al., 2011; White et al., 2012). These two traits underpin the adaptive hypotheses known as the metabolic cold adaptation (MCA) and hygric hypotheses, which aim to explain how and why metabolic rate and respiratory patterns, respectively, vary among populations and species that occupy different climatic niches (Addo-Bediako et al., 2002; Alton et al., 2017; Chown et al., 2006; Terblanche et al., 2010; White et al., 2007, 2012). However, studies that seek to understand the environmental drivers of variation in these two traits rarely consider the MCA and hygric hypotheses together, even though they are inherently linked (but see (Davis et al., 2000; Terblanche et al., 2010)).

Many insects breathe discontinuously, opening and closing their spiracles to alternate between phases of breath–holding and gas exchange with the atmosphere, and the frequency of these discontinuous gas exchange cycles increases with metabolic rate (Contreras & Bradley, 2009, 2010; Schimpf et al., 2012) (Gibbs & Johnson, 2004; White et al., 2007). A similar relationship between gas exchange frequency and metabolic rate is observed for species that breathe cyclically without closing their spiracles (Marais et al., 2005). However, whether discontinuous gas exchange patterns are adaptive remains a topic of debate (Gefen & Matthews, 2021; Gibbs & Johnson, 2004; Matthews & White, 2011; Oladipupo et al., 2022; Rowe et al., 2022; White et al., 2007).

Given the physiological link between metabolic rate and frequency of discontinuous gas exchange, it is possible that selection on either trait would result in a correlated change in the other, or a change in the relationship between them. Thus, to better understand the evolution of one trait requires consideration of the other to determine how environments have shaped trait variation and covariation. The metabolic cold adaptation hypothesis (Wohlschlag, 1960), or Krogh’s rule (Gaston et al., 2009), predicts that organisms from cold environments (such as high latitudes and altitudes) will have higher metabolic rates than those from warm environments, when measured at the same temperature. This pattern is expected to arise as a mechanism to maintain physiological function in cold environments because acute exposure to colder temperatures causes biochemical reaction rates to slow (Addo-Bediako et al., 2002; Gaston et al., 2009; Krogh, 1916; White et al., 2012). However, in terrestrial organisms, the pattern described by the MCA hypothesis may also emerge as a consequence of organisms depressing their metabolic rate in response to high temperatures to prevent desiccation (Addo– Bediako et al., 2001). This is because high air temperatures result in high vapour pressure deficits (VPDs) that can lead to higher rates of water loss from wet respiratory surfaces (White et al., 2007). Organisms living in warmer environments that are also more desiccating could therefore slow their metabolic rates, reduce their gas exchange frequency, and breathe discontinuously. The hygric hypothesis makes the explicit prediction that discontinuous gas exchange in resting tracheate arthropods, such as insects, is an adaptation to limit respiratory water loss. As such, drier environments are expected to favour arthropods that breathe discontinuously, and by extension, those that hold their breath for longer (i.e., have a lower frequency of gas exchange) (Buck et al., 1953; Chown et al., 2011; Terblanche et al., 2010; White et al., 2007).

In insects, both the MCA and hygric hypotheses tend to be supported by broad-scale comparative analyses (Addo-Bediako et al., 2002; White et al., 2007), but not always (Messamah et al., 2017). Laboratory natural selection experiments that manipulate temperature have also failed to replicate the pattern described by the MCA hypothesis in *Drosophila* (Alton et al., 2017, 2024; Mallard et al., 2018). Such research indicates that temperature alone may not be responsible for driving the evolution of metabolic rate. The hygric hypothesis similarly garners equivocal support (Chown, 2002), and alternative adaptive hypotheses propose that discontinuous gas exchange has evolved as a mechanism to improve gas exchange in underground environments, or to reduce oxidative tissue damage (Buck et al., 1953; Chown et al., 2006; Hetz & Bradley, 2005; Lighton, 1998). It has also been suggested that discontinuous gas exchange is not an adaptation to environmental conditions, but instead arises as a consequence of neural control of gas exchange during rest (Matthews & White, 2011; Rowe et al., 2022). Given that there is mixed support for both the MCA and hygric hypotheses, and that selection rarely acts on single traits independently (Blows, 2007; Calsbeek & Irschick, 2007; Lande & Arnold, 1983), we propose that these hypotheses should be considered simultaneously to better understand how metabolic rate and respiratory patterns vary and co-vary along climatic gradients.

Here we examine how metabolic rate and frequency of discontinuous gas exchange vary and co-vary within and among bee species that breathe discontinuously (da Silva et al., 2021) and exist along an altitudinal gradient on the tropical mountainous island of Viti Levu, Fiji. Altitudinal gradients are useful systems in which to test hypotheses concerning climatic drivers of trait variation because the spatial rate of change in climate is greater along altitudinal gradients than latitudinal gradients, while habitat composition and photoperiod vary less (De Frenne et al., 2013). Viti Levu is divided by a mountain range that runs north to south and the highest peak is 1328 m above sea level (a.s.l). The highlands in the centre of the island are characterised by cloud forests that are ∼5°C cooler and receive ∼100 mm more precipitation per month than the coastal lowlands (Figure 1). Due to the mountain range that divides Viti Levu, there is also a rain shadow on the western side of the island that receives significantly less rainfall than the eastern side (Figure 1c). The bee species that inhabit Viti Levu occupy different altitudinal ranges: the endemic *Homalictus fijiensis* Perkins & Cheesman 1928, is found along the entire altitudinal gradient while 22 other endemic *Homalictus* species are restricted to the highlands, and the invasive *Braunsapis puangensis* Cockerell 1928, is generally restricted to the lowlands (da Silva et al., 2016; Dorey et al., 2020; Groom et al., 2013).

**Figure 1.**
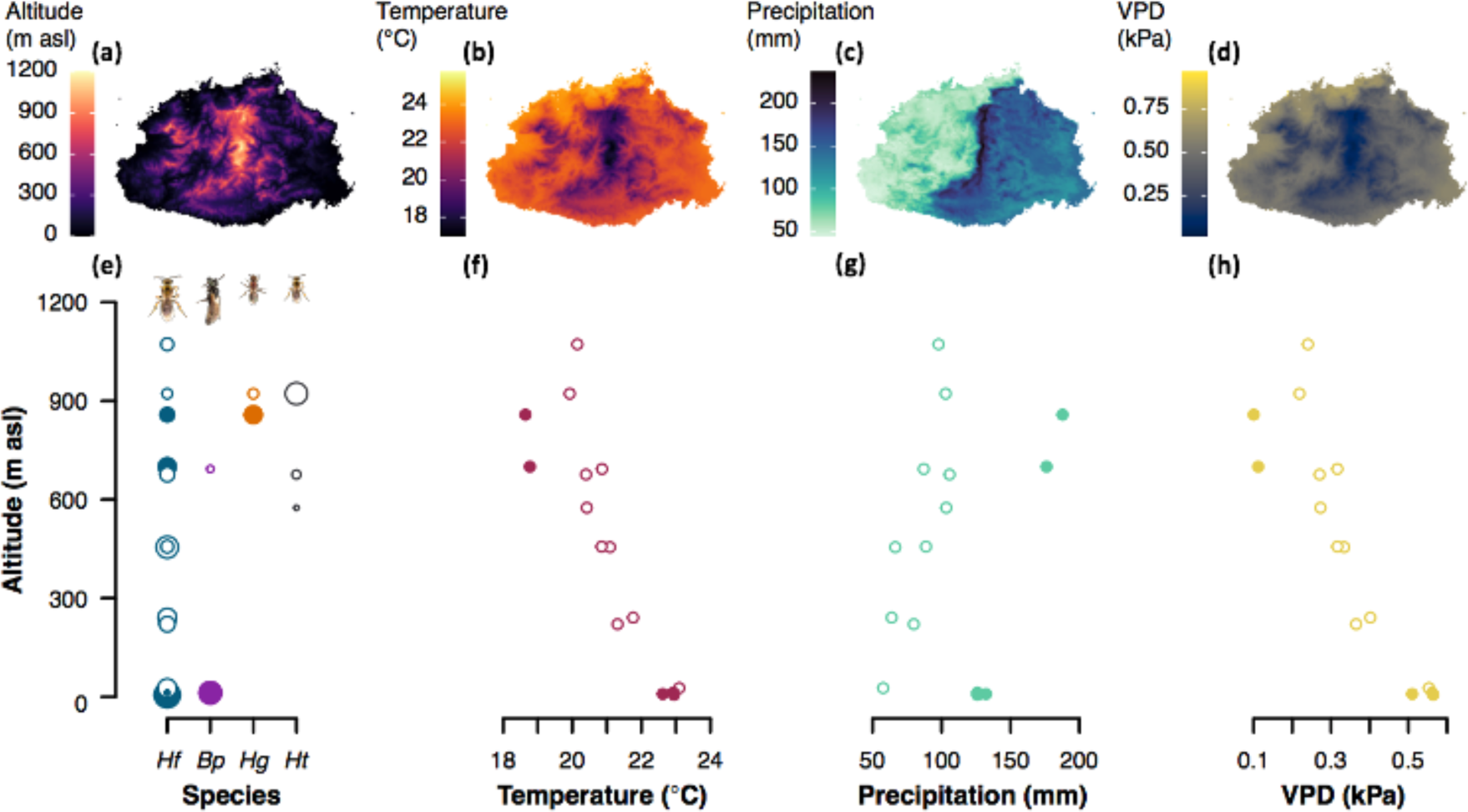
Abiotic variables across Viti Levu, Fiji: a) altitude (m asl), b) mean temperature of the coolest and driest month (July) (°C), c) precipitation of the driest month (July) (mm) and, d) vapour pressure deficit (July) (kPa). e) Collection altitude of each species (Hf: *Homalictus fijiensis*, Bp: *Braunsapis puangensis*, Hg: *Homalictus groomi*, Ht: *Homalictus tuiwawae*) where circle size indicates magnitude of sample size. f) Relationship between altitude and environmental temperature, g) relationship between altitude and precipitation, h) relationship between altitude and vapour pressure deficit (VPD). Filled circles indicate collection sites on the wet side of Viti Levu and open circles indicate sample sites on the dry side of Viti Levu.

To examine the climatic drivers of variation and co-variation in metabolic rate and respiratory patterns of bees, we collected individuals of *H. fijiensis*, *H. groomi* Dorey et al. 2019, *H. tuiwawae* Dorey et al. 2019, and *B. puangensis* from multiple sites on either side of the rain shadow along the altitudinal gradient of Viti Levu (Figure 1e). We measured the metabolic rates of 208 individual bees at 25°C (mean environmental temperature during September collection period) as rates of carbon dioxide production using flow–through respirometry. We examined the relationships between metabolic rate and frequency of gas exchange of bees and the climatic variables at their collection site using a strong inference framework whereby a set of models were selected *a priori* based on the explicit predictions of the MCA and hygric hypotheses (see Table 1 & Supplementary Box 1 for list of models tested). We accepted support for the MCA hypothesis if bees from cooler climates had higher metabolic rates and the only climate variable in the best model was temperature (Figure 2A). We accepted support for the hygric hypothesis if bees from drier climates had a lower frequency of gas exchange and the only climate variable in the best model was either vapour pressure deficit (VPD) or precipitation (Figure 2B), both of which are assumed to be indicative of aridity. In addition to testing the predictions of the MCA and hygric hypothesis, we also tested the hypothesis that frequency of gas exchange and metabolic rate are positively correlated, but that the strength of the relationship between these two traits varies depending on the aridity of the environment (Figure 2C). All else being equal, insects with higher metabolic rates breathe more frequently (Contreras & Bradley, 2009, 2010; Schimpf et al., 2012), but we expect that in drier environments there will be stronger selection for insects with high metabolic rates to maintain lower frequencies of gas exchange to minimise water loss, which will result in a shallower relationship between frequency of gas exchange and metabolic rate. In wetter environments where the risk of desiccation is lower, we expect that selection on low frequencies of gas exchange will be relaxed resulting in a steeper relationship between frequency of gas exchange and metabolic rate.

**Figure 2.**
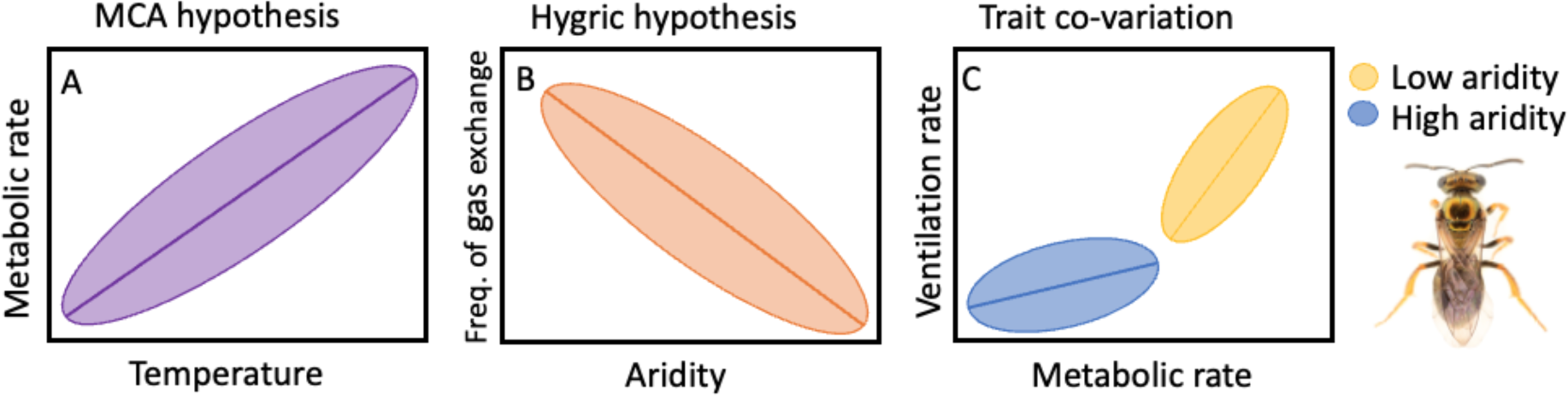
A) Conceptual illustration of the relationship between metabolic rate and temperature along an altitudinal gradient if the metabolic cold adaptation (MCA) hypothesis is supported. B) Relationship between frequency of gas exchange and aridity (inverse of precipitation or vapour pressure deficit (VPD)) if the hygric hypothesis is supported. C) Change in relationship between metabolic rate and frequency of gas exchange depending on aridity (precipitation or VPD). Where there is reduced selective pressure to avoid desiccation, we expect that selective pressure on metabolic rate and frequency of gas exchange will be relaxed, resulting in a steep relationship between metabolic rate and frequency of gas exchange in areas of high precipitation or low VPD. *Homalictus fijiensis* photograph by James B. Dorey.

**Table 1.**
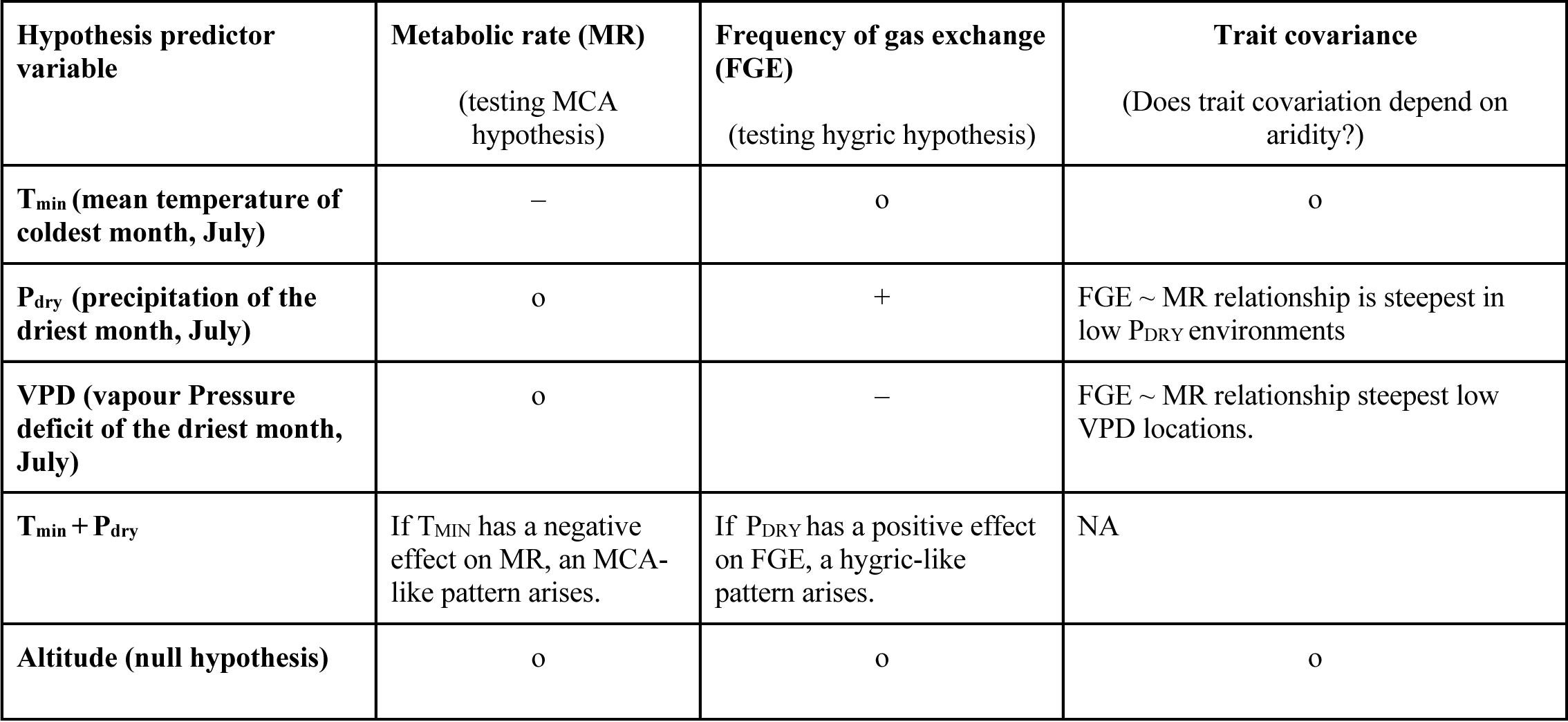
Hypotheses that were compared to test the metabolic cold adaptation hypothesis, the hygric hypothesis, and examine if metabolic rate and frequency of gas exchange (FGE) covaries depending on the climates species inhabit. A negative symbol **(–)** indicates that the hypothesis predicts a negative relationship. A positive symbol (**+)** indicates that the hypothesis predicts a positive relationship. A circle symbol (o) indicates that the hypothesis predicts a lack of a relationship between the response variable and the predictor variable.

| Hypothesis predictor variable | Metabolic rate (MR)<br>(testing MCA hypothesis) | Frequency of gas exchange (FGE)<br>(testing hygric hypothesis) | Trait covariance<br>(Does trait covariation depend on aridity?) |
| --- | --- | --- | --- |
| T<sub>min</sub> (mean temperature of coldest month, July) | – | o | o |
| P<sub>dry</sub> (precipitation of the driest month, July) | o | + | FGE ~ MR relationship is steepest in low P<sub>DRY</sub> environments |
| VPD (vapour Pressure deficit of the driest month, July) | o | – | FGE ~ MR relationship steepest low VPD locations. |
| T<sub>min</sub> + P<sub>dry</sub> | If T<sub>MIN</sub> has a negative effect on MR, an MCA-like pattern arises. | If P<sub>DRY</sub> has a positive effect on FGE, a hygric-like pattern arises. | NA |
| Altitude (null hypothesis) | o | o | o |

## Methods

### Animal collection

Bees were collected over 10 days in September 2019 on Viti Levu, Fiji. September falls within the dry season in Fiji, where precipitation and environmental temperatures are moderate (between the coolest and driest month, July, and the wettest and hottest month, February). Viti Levu is divided by a mountain range that runs north to south, which creates a rain shadow on the western side of the island that receives significantly less rainfall than the eastern side (Figure 1c). We collected bees by sweep–netting from 16 sites along an altitudinal gradient of 6–1072 m above sea level (a.s.l.), with 10 of these sites being on the dry western side of the island and the other 6 sites being on the wet eastern side. Sites were sampled in a haphazard order over the 10 days and bees were collected by sweep–netting vegetation along road verges or in native vegetation. Individuals of *Braunsapis puangensis* were also collected directly from stem nests (*Homalictus* species inhabit ground nests which we did not collect from as this is extremely laborious and yields low sample sizes). The males and females of all species collected are known to be active year–round and are multivoltine (Groom et al., 2013). Immediately upon capture, bees were placed in *Drosophila* vials with foam lids and stored in an insulated container. Inside the vials, bees were provided with a small piece of paper towel dipped in a 20% sucrose solution. On the day of collection, bees were transported to the laboratory for their metabolic rate to be measured that same evening.

### Climatic variables

For each collection site, we downloaded historical climate data (1970–2000) at a 30–arc– second resolution from WorldClim2 (Fick & Hijmans, 2017) using the R package *raster* (Hijmans et al., 2015). Since the MCA and hygric hypotheses make explicit predictions about how cool and dry climates shape metabolic rates and frequency of gas exchange, respectively, we characterised the climates at each collection site using the variables of mean temperature (°C) of the coolest month (T_min_) (July), mean precipitation (mm) of the driest month (P_dry_) (July), and mean vapour pressure (VP, kPa) of the driest month (July) (Figure 1, Table S1). VP was used to calculate the vapour pressure deficit (VPD, kPa) (drying power of the air) for bees during the driest month assuming that their respiratory surfaces are saturated with water vapour (saturated water vapour pressure, SVP, kPa) and that their body temperature matches that of the environment (T_min_, mean temperature of the driest and coolest month, °C) using the following equations:

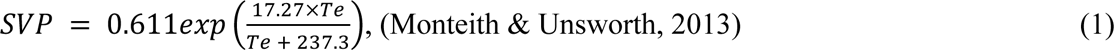

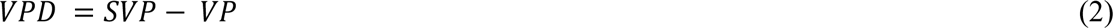

### Trait measurements

We measured the metabolic rate and frequency of gas exchange of individual bees by measuring their rates of carbon dioxide production (VCO_2_, mL h^−1^) at 25°C (mean environmental temperature across sites during collection in September) using a seven–channel flow–through respirometry system. The respirometry system was supplied with room air that was pushed through columns of soda lime and Drierite® to remove carbon dioxide and water vapour, respectively. Flow rates through each of the seven channels was regulated nominally to 100 mL min^−1^ by a mass flow controller (Aalborg, Model GFC17, Orangeburg, NY, USA). The volumetric flow rates produced by the flow controllers was measured using a Gilian Gilibrator–2 NIOSH Primary Standard Air Flow Calibrator with a low–flow cell (Sensidyne, LP, St Petersburg, FL, USA) and corrected to standard temperature pressure, STP (101.3 kPa and 0°C). To prevent bees from desiccating during measurements, the air was re–humidified after the flow controllers by passing the air through a humidifying chamber (a syringe of wetted cotton). The humidified air then flowed through a respirometry chamber (2.5 ml plastic syringe) containing an individual bee. Respirometry chambers were situated inside a temperature–controlled cabinet that maintained air temperature to 25 ± 1°C and kept bees in the dark. The excurrent air from the respirometry chamber then flowed through one of seven infrared CO_2_/H_2_O gas analysers (LI–COR, Model LI–840A, Lincoln, Nebraska, USA) that were calibrated with precision span gases (5.0 and 30.4 ppm CO_2_, Alphagaz – Air Liquide, Victoria, Australia) and measured CO_2_ concentrations at a sampling rate of 1 Hz.

The fractional CO_2_ concentration of the excurrent air (F*_eCO2_*) from each respirometry chamber was recorded for 50 min. The fractional CO_2_ concentration of the incurrent air when it was stable was measured for 2 min before and after each 50–min measurement period with an empty chamber, and a linear model was fitted to these data to estimate the CO_2_ concentration of the incurrent air during the 50–min measurement period (F*_iCO2_*). The rate of carbon dioxide production 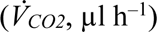 was then calculated using equation 3 (Lighton 2018) where FR is the flow rate (µl h^−1^) corrected to STP (101.3 kPa and 0°C) accounting for water vapour dilution:

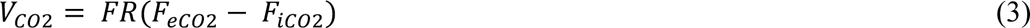

The first and last 10 min of the 60–min measurement period was considered a settling in/out period and was discarded from analysis. We measured metabolic rate over complete gas exchange cycles (from the start of the first breath until the end of the seventh breath (average measurement time = 13.6 min). Some individuals exchanged gas so slowly that 7 was the maximum number of respiratory cycles that we could measure within the 40-min measurement period. Thus, we measured metabolic rate over 7 respiratory cycles for each individual (see Supplementary Figure 1). 7 respiratory cycles is adequate for estimating metabolic rate without a large measurement error (Winwood–Smith & White, 2018).

Immediately following metabolic rate measurements, bees were preserved in 100% ethanol to be sexed and weighed at a later date. Bees were sexed by visual assessment by examining the number of antennal segments (males have 13 and females have 12). Prior to weighing, the right hind leg of each individual was detached for species identification via CO1 barcoding (see below). Dry mass was measured two months after metabolic rate measurements with specimens dried at 60°C for 48 h prior to being weighed (XP2U Ultra Micro Balance, Mettler Toledo, Greifensee, Switzerland).

Bees were measured during the evening on the day that bees were collected from the field. Due to the logistical challenges of bee collection on Viti Levu, bees could not be collected from all sites each day. Consequently, on some nights, bees from multiple sites could be measured, but on other nights only bees from one site could be measured. Since the species and sex of the bees measured each night were mostly unknown (see below), measurements were conducted blind to the species and sex of bees in most cases (*B. puangensis* is easily differentiated from *Homalictus* species). After species were identified (see below), only those species with sample sizes greater than 10 were retained for analysis. In total, we measured 208 individuals of four species: *H. fijiensis* (*n* = 125), *H. tuiwawae* (*n* = 23), *H. groomi* (*n* = 16), and *B. puangensis* (*n* = 44) (see Supplementary Table 1 for the climates that each species was collected from).

### Species identification

*Homalictus* species living at high altitudes are cryptic and, in many cases, can only be identified via male genitalia morphology, or via genetic comparison. To identify our *Homalictus* species collected from altitudes above 300 m a.s.l., we sent tissue samples (the right hind leg) to the Canadian Centre for DNA Barcoding (CCDB) at the University of Guelph, Ontario, Canada, to have the mtDNA COI gene fragment sequenced. A total of 84 samples were DNA barcoded using single–molecule real–time sequencing (SMRT49) using the PacBio Sequel platform (Pacific Biosciences, Menlo Park, CA, USA). Sequences were aligned to existing COI sequences taken from species identified by (Dorey et al., 2020, 2021). The invasive bee species, *B. puangensis*, which is generally restricted to the lowlands, is similar in size to *H. fijiensis*, but is distinguishable by its black cuticle and white facial marking (da Silva et al., 2016, 2021).

### Statistical analyses

We examined variation and co-variation in metabolic rate and frequency of gas exchange within *H. fijiensis* and among species across climatic space using linear mixed models using the nlme package (Pinheiro et al., 2017) in R Version 4.1.0 (R Development Core Team, 2019). Using a strong inference approach, we tested a suite of six models to test specific hypotheses that would support or reject the MCA and hygric hypotheses (Table 1; see rationale behind hypothesis tests in Supplementary Box 1). The sixth model in each set was a null model that included altitude and no other environmental variables (Table 1). All models included log_10_– transformed dry body mass as a predictor variable as well as variables linked to the climate hypothesis of interest. Metabolic rate was also log_10_ transformed to ensure linear mixed effect model assumptions were met. Body mass and sex were correlated, so to avoid collinearity issues, we elected to remove sex as a predictor factor from our models as it is crucial to include body mass when examining variation in metabolic rate (Lighton, 2018). To deal with collinearity between the environmental variables in our models (and as a result, high variance inflation factors, VIFs), especially between temperature, VPD, and altitude, we modelled each environmental variable separately, except for a model that included temperature and precipitation of the driest month as VIFs were under 2 (Johnston et al., 2018). Models testing the hygric hypothesis included log_10_-transformed metabolic rate as a predictor variable to account for the effect of metabolic rate on frequency of gas exchange as they are positively correlated. Multispecies models testing the MCA and the hygric hypothesis included species as a fixed factor within the models. Measurement block (sets of bees measured at the same time) and channel were included as random intercepts within each model.

We compared the Akaike Information Criterion (AIC) value of each model to determine which hypothesis best explained the variation in metabolic rate, frequency of gas exchange and how traits co–vary across abiotic variables as per Burnham & Anderson (2002) and White et al., (2007) using the MuMIn package (Barton & Barton, 2015). We then evaluated the direction of the relationships between the traits of interest and abiotic variables within the best fitting models to make inferences on whether metabolic rate and frequency of gas exchange are best explained by the MCA and hygric hypotheses, respectively, and if the relationship between metabolic rate and frequency of gas exchange remains consistent across abiotic variables or if it becomes uncoupled. All models included log_10_–transformed body mass as a predictor variable, and an interaction between log_10_–transformed body mass and species were included within the multispecies models. The multispecies models testing the hygric hypothesis also included an interaction between log_10_–transformed metabolic rate and species. Only species that had sample sizes over 10 individuals were included in the multispecies analysis (four species). The packages ggplot2 (Wickham, 2011) and cowplot (Wilke et al., 2019) were used to produce data figures throughout the results.

To examine if the relationship (regression slope) between metabolic rate and frequency of gas exchange depends on the climatic conditions species inhabit, we attempted to run a series of linear mixed effect models. Frequency of gas exchange was the response variable, log_10_– transformed body mass was a fixed–factor predictor variable, and a three-way interaction between the fixed factors of species, metabolic rate and each climatic factor (each climatic factor would be modelled in a separate model as per above) would inform on whether the slope changes depending on the different environments species inhabit. However, because sample sizes were small for our two highland species (n = 16 and n = 23, for *H. groomi* and *H. tuiwawae,* respectively) and *H. groomi* was collected in a very narrow elevational band (two sites), we were underpowered to adequately test this three–way interaction. However, we were able to estimate the slope of relationship between frequency of gas exchange and metabolic rate for each species while accounting for body mass by running separate linear mixed effect models for each species. Frequency of gas exchange was the response variable, and log_10_– transformed metabolic rate and log_10_–transformed body mass were the fixed–factor predictor variables, and measurement block and channel were included as random intercepts. We then calculated the mean collection altitude, mean temperature of the coldest month, mean precipitation of the driest month, and mean VPD, that each species inhabits across the elevational gradient. We used linear models where slope of the relationship between frequency of gas exchange and metabolic rate was the response variable, and each climatic variable was the fixed-factor predictor variable to evaluate whether the slope of the relationship between frequency of gas exchange and metabolic rates changes depends on aridity. The five models were compared using AIC as per above.

## Results

### Testing the MCA within and among species

The model that best explained variation in metabolic rate within *H. fijiensis* and across species included average temperature of the coldest month (*H. fijiensis*: estimate = –0.041 ± SE = 0.012, df = 92, t = –3.3, P < 0.001) (among species: (estimate = –0.037 ± SE = 0.013, df = 167, t = –2.9, P = 0.004), and average precipitation of the driest month as the test variables (*H. fijiensis*: estimate = –0.002 ± SE = 0.0005, df = 92, t = –3.6, P < 0.005) (among species: estimate = –0.002 ± SE = 0.0004, df = 167, t = –3.44, P < 0.005) (Table 2 and see full model summaries in Supplementary Tables 2 and 3). Metabolic rate was negatively correlated with both temperature and precipitation, therefore an MCA-like pattern was observed (Figure 3A & 2B), but the MCA hypothesis was not supported in its strictest form as precipitation also explained a proportion of the variation in metabolic rate. Body mass was positively correlated with metabolic rate within *H. fijiensis* (estimate = 0.76 ± SE = 0.07, df = 92, t = 11.6, P < 0.001) and across bee species (estimate = 0.74 ± SE = 0.06, df = 167, t = 12.37, P < 0.001).

**Table 2.**
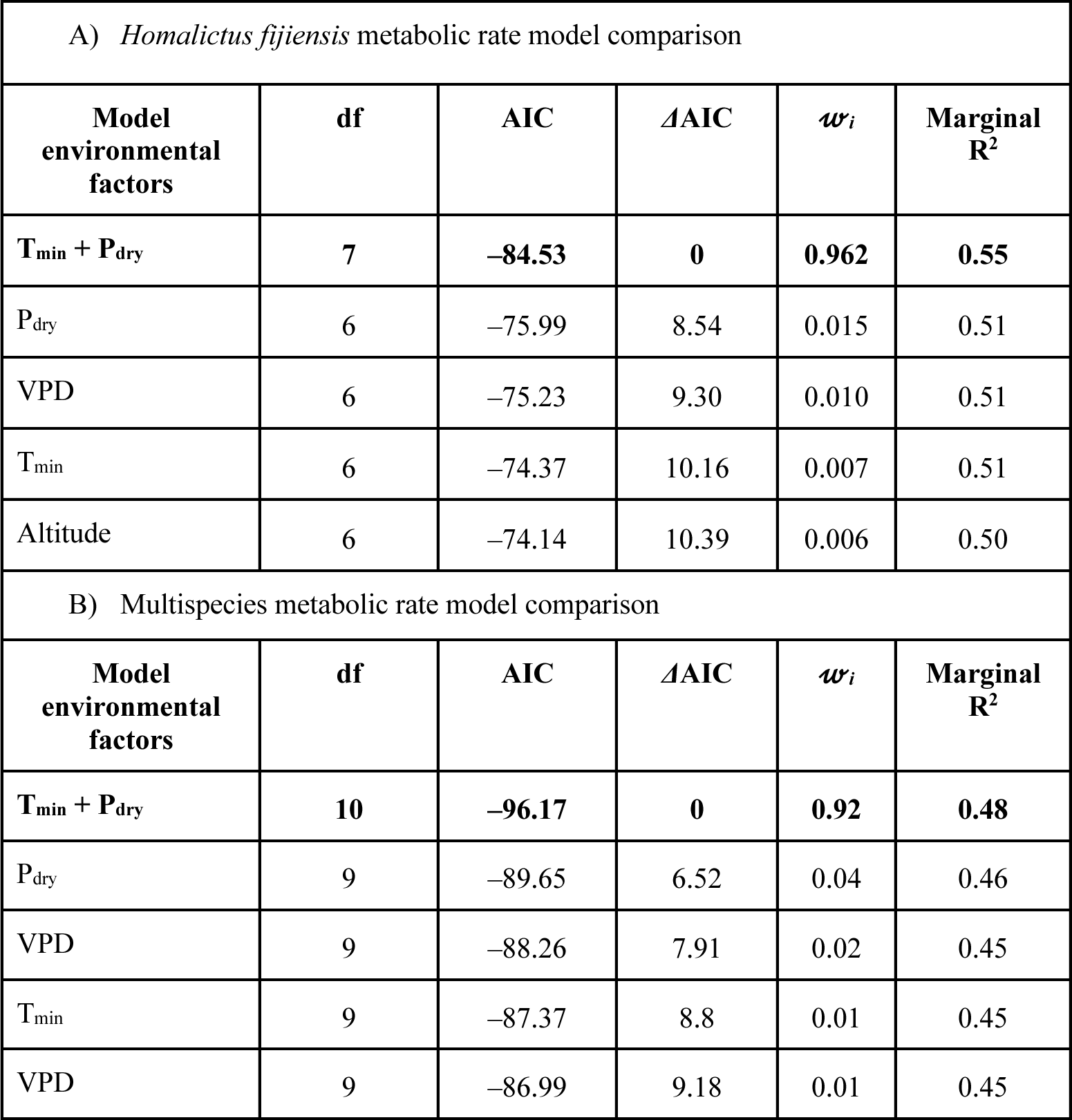
Testing the MCA within *H. fijiensis* (A) and among species (B). Model comparison table. Best fitting model with lowest AIC value is in bold. T_min_ = mean environmental temperature of coldest month (°C), P_dry_= mean precipitation of the driest month (mm), VPD = vapour pressure deficit (kPa). All models also included log_10_–transformed body mass as predictor variable.*Wi* indicates AIC model weight.

| A) *Homalictus fijiensis* metabolic rate model comparison |  |  |  |  |  |
| --- | --- | --- | --- | --- | --- |
| Model environmental factors | df | AIC | $\Delta\text{AIC}$ | $w_i$ | Marginal $R^2$ |
| **$T_{\min} + P_{\text{dry}}$** | **7** | **-84.53** | **0** | **0.962** | **0.55** |
| $P_{\text{dry}}$ | 6 | -75.99 | 8.54 | 0.015 | 0.51 |
| VPD | 6 | -75.23 | 9.30 | 0.010 | 0.51 |
| $T_{\min}$ | 6 | -74.37 | 10.16 | 0.007 | 0.51 |
| Altitude | 6 | -74.14 | 10.39 | 0.006 | 0.50 |
| B) Multispecies metabolic rate model comparison |  |  |  |  |  |
| Model environmental factors | df | AIC | $\Delta\text{AIC}$ | $w_i$ | Marginal $R^2$ |
| **$T_{\min} + P_{\text{dry}}$** | **10** | **-96.17** | **0** | **0.92** | **0.48** |
| $P_{\text{dry}}$ | 9 | -89.65 | 6.52 | 0.04 | 0.46 |
| VPD | 9 | -88.26 | 7.91 | 0.02 | 0.45 |
| $T_{\min}$ | 9 | -87.37 | 8.8 | 0.01 | 0.45 |
| VPD | 9 | -86.99 | 9.18 | 0.01 | 0.45 |

**Figure 3.**
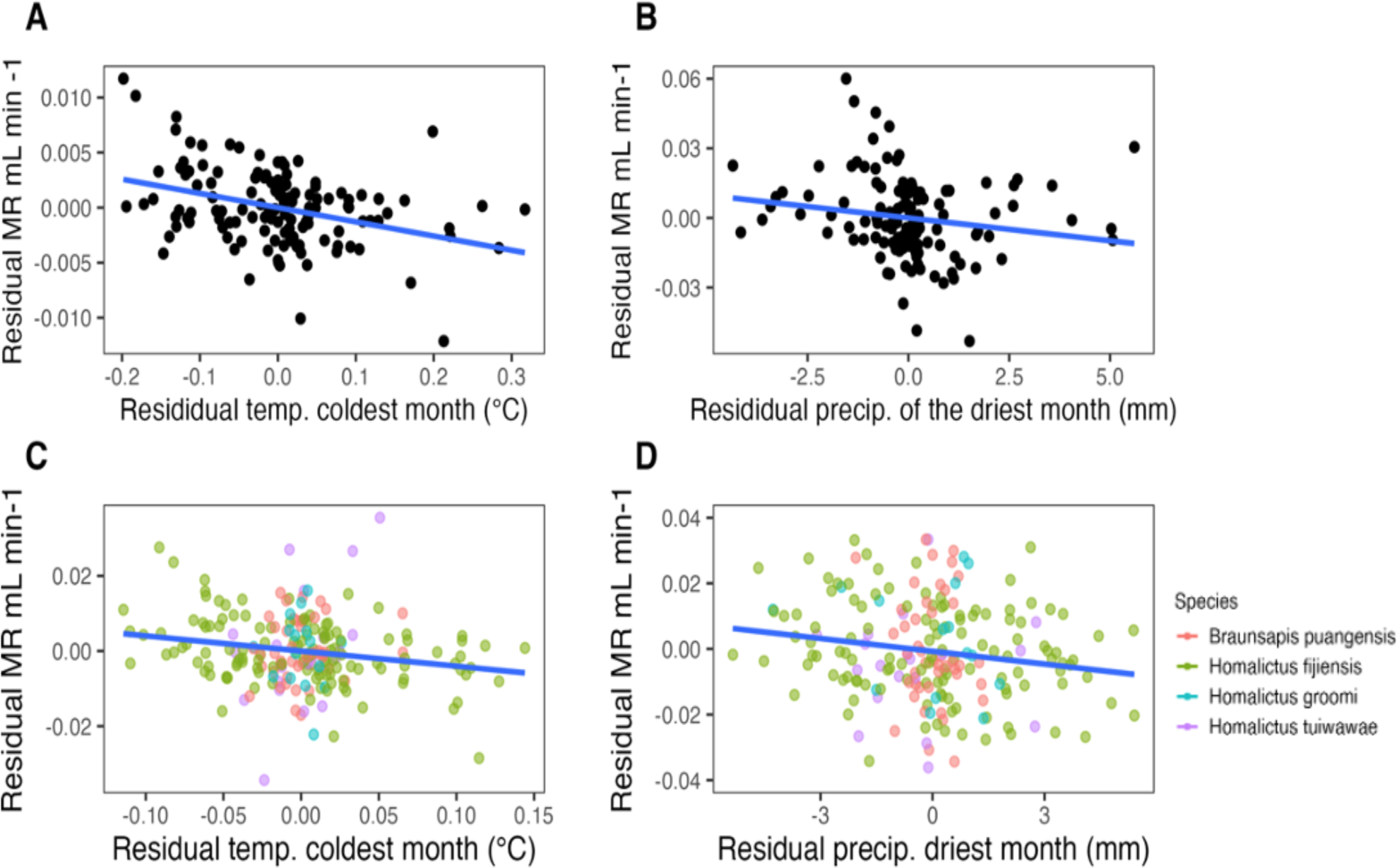
A) Residual metabolic rate (mL hr^−1^) of *H. fijiensis* across residual average temperature of the coldest month (accounting for variation in body size), and B) residual precipitation of the driest month (mm). C) Residual metabolic rate of multiple Fijian bee species (accounting for variation in body mass) across residual temperature of the coldest month, and D) residual precipitation of the driest month. Note that residuals are presented for data visualisation purposes as per White et al., (2007), however, the raw data was analysed.

### Testing the hygric hypothesis within and across species

The model that best explained variation in frequency of gas exchange within *H. fijiensis and* across species included only altitude as the explanatory environmental variable (Table 3). However, altitude did not have a significant effect on frequency of respiration in *H. fijiensis* or among species (see Supplementary Tables 4 and 5 for full model coefficient summaries). The best model was only marginally more supported than models that included precipitation of the driest month and temperature or precipitation of the driest month as the only environmental variable within the *H. fijiensis* models, and models that included precipitation of the driest month, temperature, and VPD in the among species models, based on the ΔAIC values (< 2 ΔAIC). However, none of the variables in the models that performed marginally worse than the best model were significantly correlated with frequency of gas exchange in either *H. fijiensis* or among species. Thus, the hygric hypothesis was not supported. Body mass was negatively correlated with frequency of gas exchange within *H. fijiensis* (estimate = –0.51 ± SE = 0.08, df = 92, t = –5.75, P < 0.001) and among bee species (estimate = –0.43 ± SE = 0.07, df = 167, t = –5.96, P < 0.001) (i.e. smaller bees have more rapid gas exchange frequencies than larger bees). Frequency of respiration was positively correlated with metabolic rate within *H. fijiensis* (estimate = 0.67 ± SE = 0.09, df = 92, t = 7.74, P < 0.001) and among species (estimate = 0.55 ± SE = 0.06, df = 167, t = 9.05, P < 0.001) (Figure 4).

**Table 3.**
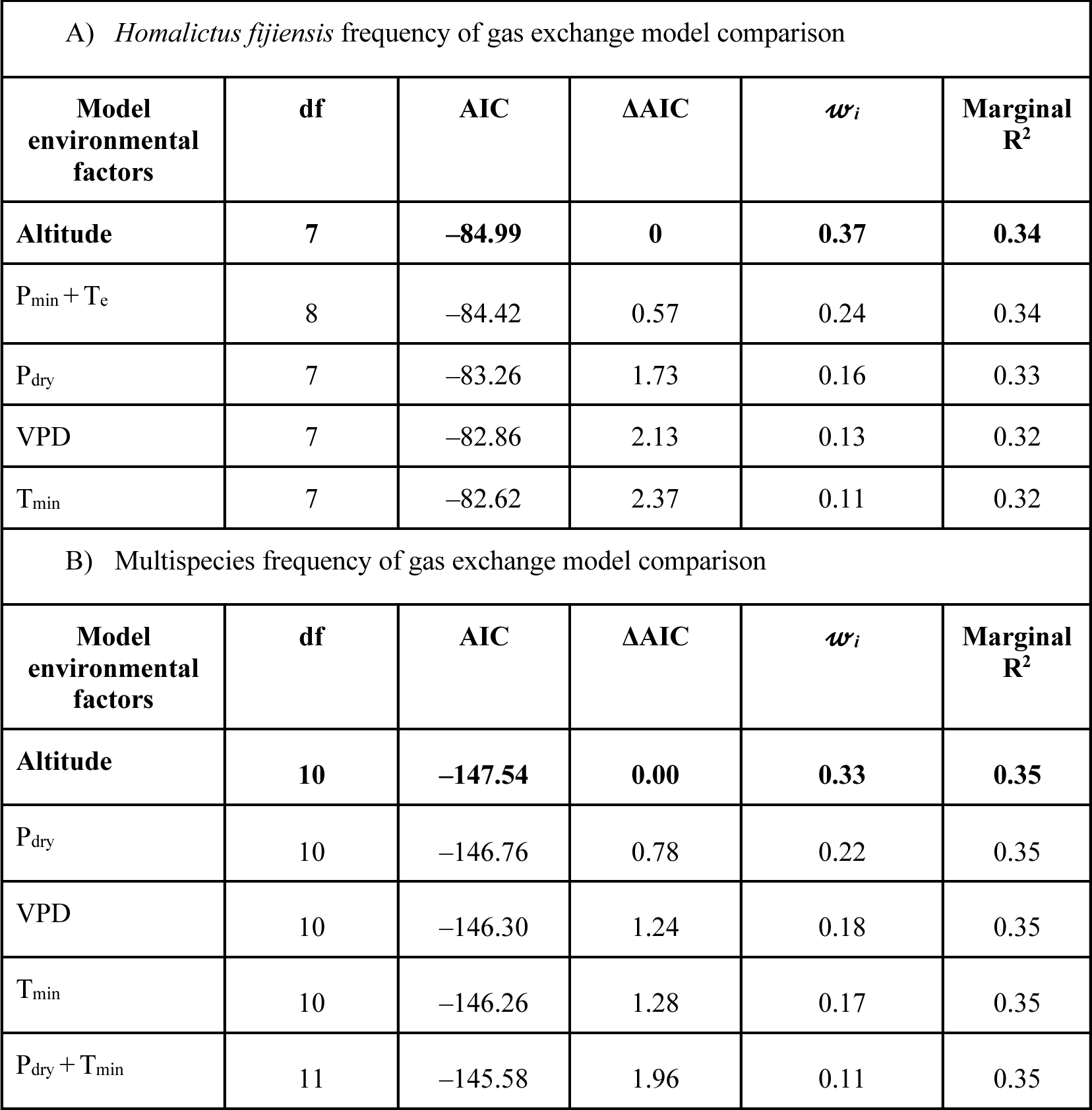
Testing the hygric hypothesis within *Homalictus fijiensis* (A) and across Fijian bee species (B). Best fitting model with lowest AIC value is in bold. T_min_ = mean environmental temperature of coldest month (°C), P_dry_ = mean precipitation of the driest month (mm), VPD = vapour pressure deficit (kPa). All models also included log_10_–transformed body mass, altitude and log_10_–transformed metabolic rate as fixed–factor predictor variables.*W_i_* indicates model <u>weight.</u>

**Figure 4.**
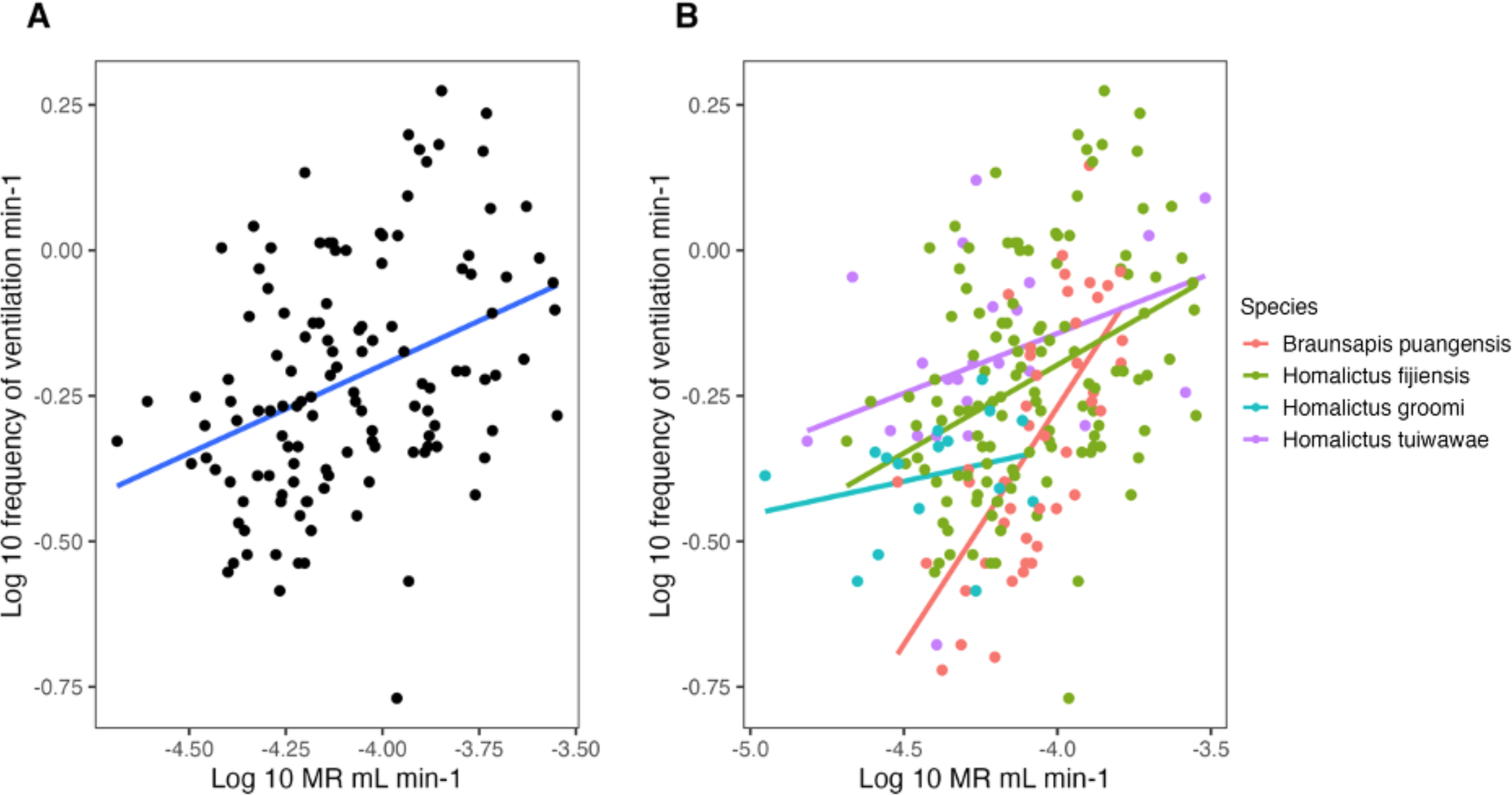
Relationship between the frequency of gas exchange (frequency of gas exchange min^−1^) and metabolic rate (mL hr^−1^) within A) *H. fijiensis*, and B) among species.

### Does the relationship between metabolic rate and frequency of gas exchange co–vary depending on the different climates species inhabit?

The model that included mean collection altitude best explained variation in the slope of the relationship between frequency of gas exchange and metabolic rate across species (R^2^ = 0.97) (Table 4) (Supplementary Table 6). The relationship between frequency of gas exchange and metabolic rate is steeper in species that occupy low elevation regions (*B. puangensis* and *H. fijiensis*) compared to species that are restricted to high elevations (*H. groomi* and *H. tuiwawae*) (Figure 5).

**Table 4.** Trait covariation model comparison. Best fitting model with lowest AIC value is in bold. T_min_ = mean environmental temperature of coldest month (°C), P_dry_= mean precipitation of the driest month (mm), VPD = vapour pressure deficit (kPa).

| A) Multispecies trait covariance model comparison (response variable is the slope of the relationship between frequency of gas exchange and metabolic rate) |  |  |  |  |
| --- | --- | --- | --- | --- |
| Testing how trait covariance changes with climatic variables | df | AIC | $\Delta\text{AIC}$ | $R^2$ |
| **Altitude** | **3** | **-6.05** | **0.00** | **0.97** |
| VPD | 3 | -0.46 | 5.59 | 0.87 |
| $T_{\min}$ | 3 | -0.11 | 5.94 | 0.86 |
| $T_{\min} + P_{\text{dry}}$ | 4 | 0.98 | 7.03 | 0.89 |
| $P_{\text{dry}}$ | 3 | 7.49 | 13.54 | 0.08 |

**Figure 5.**
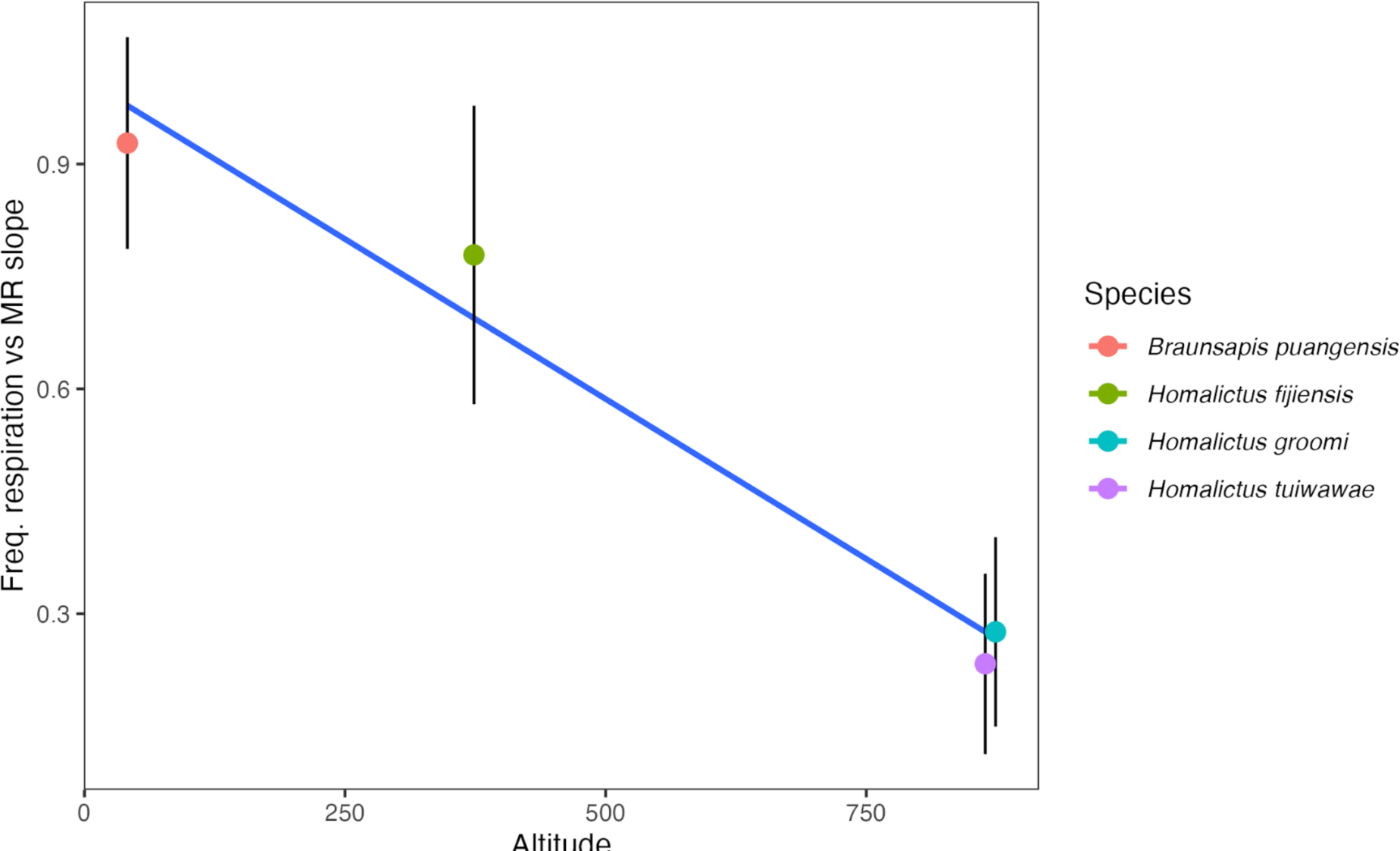
Slope of the relationship (accounting for body mass) between frequency of gas exchange and metabolic rate depending on the mean elevation species were collected from (R^2^ = 0.97).

## Discussion

Adaptive hypotheses that explain variation in physiological traits along climatic gradients make explicit predictions based on the expected direct effects of abiotic variables on physiological function. The metabolic cold adaptation (MCA) and hygric hypotheses predict that metabolic rates and gas exchange patterns evolve in response to the effects of temperature and water availability on these two traits, respectively (Addo-Bediako et al., 2002; White et al., 2007). Whether these hypotheses are supported within and among species and across environments has been the centre of debate over the last few decades (Addo-Bediako et al., 2002; Alton et al., 2024; Clarke, 2003; Messamah et al., 2017; Oladipupo et al., 2022; Rowe et al., 2022; White et al., 2007). However, despite metabolic rate and gas exchange patterns being inherently linked, how they covary across climatic conditions is rarely considered. Understanding how climate shapes traits and their relationships is likely to be critical for improving our understanding of the mechanisms that underpin species distributions and survival with further climate change (Hellmann & Pineda–Krch, 2007; Svensson et al., 2021). We discuss how metabolic rate and frequency of gas exchange vary across climatic gradients independently and then simultaneously to explore how these traits might evolve together.

We found that the MCA hypothesis was not supported in its strictest form in Fijian bees. Although we found a negative relationship between metabolic rate and temperature, as predicted by the MCA hypothesis, the best model that explained variation in metabolic rate also included precipitation. We expected that if precipitation did explain variation in metabolic rate, that metabolic rate would be lowest in dry environments and highest in wet environments as a mechanism to avoid desiccation (Supplementary Box 1). However, we found that metabolic rate was negatively correlated with precipitation (Figure 3B & 3D). This pattern might be explained by variation in environmental productivity and competition. Productivity is usually greatest in regions of high precipitation, and biodiversity is often positively correlated with productivity (Fei et al., 2018). Land surface area decreases with altitude in Fiji, leading to an environment that is small, but productive, and contains high endemic bee diversity (over 22 species in highlands) (Dorey et al., 2020). Therefore, high competition for resources between species could lead to a situation where lack of energetic resources (due to high competition despite high productivity) could be reducing metabolic rates in bees that inhabit high–precipitation, high–altitude regions. Low metabolic rates might be favoured in environments with high competition or fewer resources following the ‘compensation’ hypothesis which suggests that individuals with low resting metabolic rates will have higher fitness because they have lower self-maintenance costs and can devote more energy towards growth and reproduction (Burton et al., 2011). However, this hypothesis is contradicted by recent research which shows that individuals with higher metabolic rates have greater fitness when intraspecific competition is intense (Pettersen et al., 2020), and that competition leads to higher metabolic rates (in *Drosophila*) due to increased activity (at high temperatures) (Alton & Kellermann, 2023).

Variation in the frequency of gas exchange within *H. fijiensis* and across Fijian bee species was not significantly explained by any environmental variables included in our analysis (Table 3). Therefore, we do not find any empirical support for the hygric hypothesis within bees in Fiji. Discontinuous gas exchange has evolved independently at least five times (Marais et al., 2005), and thus, there could be multiple mechanisms that explain why discontinuous gas exchange occurs in insects, and the variation in frequency of gas exchange we observe within species that breathe discontinuously (Chown, 2002; Matthews & White, 2011). The chthonic hypothesis (Lighton, 1998) posits that breathing discontinuously improves gas exchange in underground (high CO_2_) environments. Although this hypothesis, doesn’t tend to garner much empirical support (Chown & Holter, 2000; White et al., 2007), including within a recent meta-analysis (Oladipupo et al., 2022), it could explain the evolution of discontinuous gas exchange in bees. *Homalictus* bee species nest in deep (often over 1 m deep) ground nests in large groups (Danforth & Ji, 2001), which could potentially create high CO_2_ environments. *B. puangensis* nest in stems above the ground, but many individuals will often live together within short linear nests (da Silva et al., 2016), and they are known to plug their entrance holes with their abdomens to keep rain and predators out of the nest (pers obs. da Silva & Schwarz 2019; and a known behaviour in other allodapine bee species (Melna & Schwarz, 1994)). Therefore, *B. puangensis* might also live in high CO_2_ nest environments, despite living above ground.

Alternatively, discontinuous gas exchange could be a conserved state across all bee species when they are in a state of rest. Matthews and White (2011) proposed that discontinuous gas exchange occurs as a consequence of reduced or down regulated brain activity. *Homalictus spp*. and *B. puangensis* belong to different families, Halictidae and Apidae, respectively, and they both exhibited discontinuous gas exchange when placed in the dark metabolic rate chamber, as do other species within Apidae (Lighton & Lovegrove, 1990; Beekman & Van Stratum, 1999) and Megachilidae (Grula et al., 2021). Thus, perhaps variation in the frequency of gas exchange within and among species can be attributed to the degree of brain activity while at rest (Matthews & White, 2011), as well as metabolic rate and body mass (this study & Terblanche et al., 2008). Finally, the oxidative damage hypothesis (Bradley, 2000), suggests that discontinuous gas exchange evolves to reduce toxic effects of near ambient intratracheal oxygen levels. Unfortunately, we have no way of evaluating whether this hypothesis could explain variation in the frequency of gas exchange in the current study and thus cannot speculate on whether our data might support it or not. However, the oxidative damage hypothesis seems to have very little empirical support by other studies (Matthews & White, 2011; White et al., 2007).

The relationship between frequency of gas exchange and metabolic rate changed across species in our study depending on their mean elevational range. The two species which dominate the lowland region, *B. puangensis* and *H. fijiensis*, had a much steeper relationship, where frequency of gas exchange increased more quickly with metabolic rate, than in the two highland species, *H. groomi* and *H. tuiwawae*. While the model that included altitude as the environmental factor had the lowest AIC value (see Table 4) and the highest R^2^ value (0.97), models that included VPD (R^2^ = 0.87), environmental temperature (R^2^ = 0.86), and environmental temperature and precipitation of the driest month (R^2^ = 0.89) also explained a large proportion of the variation in the slope of the relationship between frequency of gas exchange and metabolic rate. This suggests that temperature, aridity, and another factor associated with elevation that we did not test could explain how frequency of gas exchange and metabolic rate co-vary across the environmental gradient. For example, perhaps elevation is correlated with nest depth or complexity, which could contribute towards nest CO_2_ levels if the chthonic hypothesis explains frequency of gas exchange.

While frequency of gas exchange and metabolic rate are linked (they are positively correlated across all species) we could be observing a physiological trait trade-off. While we can not determine whether the change in correlation between traits across the elevational gradient is due to correlational selection or a genetic correlation in this study, the fact that species from different environments have different trait relationships suggests that there is scope for metabolic rate and frequency of gas exchange to evolve independently of each other depending on environmental or other ecological factors. However, while the two traits might be evolving independently of each other, they remain correlated, and the extent to which each trait can evolve independently in response to variation in climate remains an exciting research question for the future.

We believe it is critical to examine organisms as a mosaic of traits to improve our understanding of organismal responses to changing environments (Endler, 1995; Roff & Fairbairn, 2012; Svensson et al., 2021). Understanding whether key adaptive hypotheses, such as the MCA, are supported across species is crucial for predicting how climate change will impact species energetics in the future. However, variation in the strength of trait relationships (such as metabolic rate and frequency of gas exchange) could have implications for the ways in which species evolve with further anthropogenic climate change (Chantepie & Chevin, 2020). As organisms are composed of multiple interacting traits and adaptation is a multifactorial process, we believe it is important for future research to consider how multiple traits evolve across landscapes together as we move further into the Anthropocene.

## Acknowledgements

We would like to thank Craig White for access to equipment, helpful discussions and providing comments on the manuscript. We also thank James Dorey for providing bee photographs. We would also like to thank the Navai village for hosting us. This research was funded by an Endeavour Leadership Postdoctoral Scholarship (7185_2019), a Company of Biologist Travelling Fellowship and a Macquarie University Research Fellowship to CRBdS. Funding from Monash University (Advancing Women in Science Grant) was awarded to VK. This research was supported by funding from the Australian Research Council (DP180103925 and DP220103421 to LAA, DP200101272 to VK). This research was also partially funded by the Australian Department of Foreign Affairs and Trade (via the New Colombo Plan program) to MPS and MIS. The study was carried out in Fiji under permit number 1755–11.

## Author contributions

CRBdS, JEB, VK, and LAA conceptualised the manuscript. CRBdS and JEB conducted the metabolic rate experiments. MT, MIS, and MPS provided helpful logistical support in Fiji. CRBdS extracted the data, conducted the statistical analyses, and wrote the first draft of the manuscript. All authors edited the manuscript.

## Data availability statement

Data will be published on Dryad upon manuscript acceptance.

## References

Addo–Bediako, A., Chown, S. L., & Gaston, K. J. (2001). Revisiting water loss in insects: A large scale view. Journal of Insect Physiology, 47(12), 1377–1388.

Addo-Bediako, A., Chown, S. L., & Gaston, K. J. (2002). Metabolic cold adaptation in insects: A large-scale perspective. Functional Ecology, 16(3), 332–338.

Alton, L. A., Condon, C., White, C. R., & Angilletta Jr, M. J. (2017). Colder environments did not select for a faster metabolism during experimental evolution of *Drosophila melanogaster*. Evolution, 71(1), 145–152.

Alton, L. A., & Kellermann, V. (2023). Interspecific interactions alter the metabolic costs of climate warming. Nature Climate Change, 13(4), 382–388.

Alton, L. A., Kutz, T., Bywater, C. L., Lombardi, E., Cockerell, F. E., Layh, S., Winwood–Smith, H., Arnold, P. A., Beaman, J. E., & Walter, G. M. (2024). Temperature and nutrition do not interact to shape the evolution of metabolic rate. Philosophical Transactions of the Royal Society B, 379(1896), 20220484.

Angilletta Jr, M. J., Huey, R. B., & Frazier, M. R. (2010). Thermodynamic effects on organismal performance: Is hotter better? Physiological and Biochemical Zoology, 83(2), 197–206.

Barton, K., & Barton, M. K. (2015). Package ‘mumin.’ Version, 1(18), 439.

Beekman, M., & Van Stratum, P. (1999). Respiration in bumblebee queens: Effect of life phase on the discontinuous ventilation cycle. Entomologia Experimentalis et Applicata, 92(3), 295–298.

Blows, M. W. (2007). A tale of two matrices: Multivariate approaches in evolutionary biology. Journal of Evolutionary Biology, 20(1), 1–8.

Bradley, T. J. (2000). The discontinuous gas exchange cycle in insects may serve to reduce oxygen supply to the tissues. American Zoologist, 40(6), 952–952.

Buck, J., Keister, M., & Specht, H. (1953). Discontinuous respiration in diapausing Agapema pupae. Anat. Rec, 117(541), 541.

Burnham, K. P., & Anderson, D. R. (2002). A practical information–theoretic approach. Model Selection and Multimodel Inference, 2.

Burton, T., Killen, S. S., Armstrong, J. D., & Metcalfe, N. B. (2011). What causes intraspecific variation in resting metabolic rate and what are its ecological consequences? Proceedings of the Royal Society B: Biological Sciences, 278(1724), 3465–3473.

Calsbeek, R., & Irschick, D. J. (2007). The quick and the dead: Correlational selection on morphology, performance, and habitat use in island lizards. Evolution: International Journal of Organic Evolution, 61(11), 2493–2503.

Chantepie, S., & Chevin, L. (2020). How does the strength of selection influence genetic correlations? Evolution Letters, 4(6), 468–478.

Chown, S. L. (2002). Respiratory water loss in insects. Comparative Biochemistry and Physiology Part A: Molecular & Integrative Physiology, 133(3), 791–804.

Chown, S. L., Gibbs, A. G., Hetz, S. K., Klok, C. J., Lighton, J. R., & Marais, E. (2006). Discontinuous gas exchange in insects: A clarification of hypotheses and approaches. Physiological and Biochemical Zoology, 79(2), 333–343.

Chown, S. L., & Holter, P. (2000). Discontinuous gas exchange cycles in *Aphodius fossor* (Scarabaeidae): A test of hypotheses concerning origins and mechanisms. Journal of Experimental Biology, 203(2), 397–403.

Chown, S. L., Sørensen, J. G., & Terblanche, J. S. (2011). Water loss in insects: An environmental change perspective. Journal of Insect Physiology, 57(8), 1070–1084.

Clarke, A. (2003). Costs and consequences of evolutionary temperature adaptation. Trends in Ecology & Evolution, 18(11), 573–581.

Contreras, H. L., & Bradley, T. J. (2009). Metabolic rate controls respiratory pattern in insects. Journal of Experimental Biology, 212(3), 424–428.

Contreras, H. L., & Bradley, T. J. (2010). Transitions in insect respiratory patterns are controlled by changes in metabolic rate. Journal of Insect Physiology, 56(5), 522– 528.

da Silva, C. R. B., Stevens, M. I., & Schwarz, M. P. (2016). Casteless sociality in an allodapine bee and evolutionary losses of social hierarchies. Insectes Sociaux, 63(1), 67–78.

da Silva, C. R., Beaman, J. E., Dorey, J. B., Barker, S. J., Congedi, N. C., Elmer, M. C., Galvin, S., Tuiwawa, M., Stevens, M. I., & Alton, L. A. (2021). Climate change and invasive species: A physiological performance comparison of invasive and endemic bees in Fiji. Journal of Experimental Biology, 224(1), jeb230326.

da Silva, C. R. B., Groom, S. V., Stevens, M. I., & Schwarz, M. P. (2016). Current status of the introduced allodapine bee *Braunsapis puangensis* (Hymenoptera: Apidae) in Fiji. Austral Entomology, 55(1), 43–48.

Danforth, B. N., & Ji, S. (2001). Australian *Lasioglossum*+ *Homalictus* form a monophyletic group: Resolving the “Australian enigma.” Systematic Biology, 50(2), 268–283.

Davis, A. L., Chown, S. L., McGeoch, M. A., & Scholtz, C. H. (2000). A comparative analysis of metabolic rate in six *Scarabaeus* species (Coleoptera: Scarabaeidae) from southern Africa: Further caveats when inferring adaptation. Journal of Insect Physiology, 46(4), 553–562.

De Frenne, P., Graae, B. J., Rodríguez-Sánchez, F., Kolb, A., Chabrerie, O., Decocq, G., De Kort, H., De Schrijver, A., Diekmann, M., & Eriksson, O. (2013). Latitudinal gradients as natural laboratories to infer species’ responses to temperature. Journal of Ecology, 101(3), 784–795.

Diamond, S. E., & Martin, R. A. (2021). Physiological adaptation to cities as a proxy to forecast global–scale responses to climate change. Journal of Experimental Biology, 224(Suppl_1), jeb229336.

Dorey, J. B., Groom, S. V., Freedman, E. H., Matthews, C. S., Davies, O. K., Deans, E. J., Rebola, C., Stevens, M. I., Lee, M. S., & Schwarz, M. P. (2020). Radiation of tropical island bees and the role of phylogenetic niche conservatism as an important driver of biodiversity. Proceedings of the Royal Society B, 287(1925), 20200045.

Dorey, J. B., Groom, S. V., Velasco-Castrillón, A., Stevens, M. I., Lee, M. S., & Schwarz, M. P. (2021). Holocene population expansion of a tropical bee coincides with early human colonization of Fiji rather than climate change. Molecular Ecology, 30(16), 4005–4022.

Endler, J. A. (1995). Multiple–trait coevolution and environmental gradients in guppies. Trends in Ecology & Evolution, 10(1), 22–29.

Fei, S., Jo, I., Guo, Q., Wardle, D. A., Fang, J., Chen, A., Oswalt, C. M., & Brockerhoff, E. G. (2018). Impacts of climate on the biodiversity–productivity relationship in natural forests. Nature Communications, 9(1), 1–7.

Fick, S. E., & Hijmans, R. J. (2017). WorldClim 2: New 1-km spatial resolution climate surfaces for global land areas. International Journal of Climatology, 37(12), 4302– 4315.

Gaston, K. J., Chown, S. L., Calosi, P., Bernardo, J., Bilton, D. T., Clarke, A., Clusella–Trullas, S., Ghalambor, C. K., Konarzewski, M., & Peck, L. S. (2009). Macrophysiology: A conceptual reunification. The American Naturalist, 174(5), 595– 612.

Gefen, E., & Matthews, P. G. (2021). From chemoreception to regulation: Filling the gaps in understanding how insects control gas exchange. Current Opinion in Insect Science, 48, 26–31.

Gibbs, A. G., & Johnson, R. A. (2004). The role of discontinuous gas exchange in insects: The chthonic hypothesis does not hold water. Journal of Experimental Biology, 207(20), 3477–3482.

Groom, S. V., Stevens, M. I., & Schwarz, M. P. (2013). Diversification of Fijian halictine bees: Insights into a recent island radiation. Molecular Phylogenetics and Evolution, 68(3), 582–594.

Grula, C. C., Rinehart, J. P., Greenlee, K. J., & Bowsher, J. H. (2021). Body size allometry impacts flight–related morphology and metabolic rates in the solitary bee *Megachile rotundata*. Journal of Insect Physiology, 133, 104275.

Hellmann, J. J., & Pineda–Krch, M. (2007). Constraints and reinforcement on adaptation under climate change: Selection of genetically correlated traits. Biological Conservation, 137(4), 599–609.

Hetz, S. K., & Bradley, T. J. (2005). Insects breathe discontinuously to avoid oxygen toxicity. Nature, 433(7025), 516–519.

Hijmans, R. J., Van Etten, J., Cheng, J., Mattiuzzi, M., Sumner, M., Greenberg, J. A., Lamigueiro, O. P., Bevan, A., Racine, E. B., & Shortridge, A. (2015). Package ‘raster.’ R Package, 734.

Hoffmann, A. A., & Sgro, C. M. (2011). Climate change and evolutionary adaptation. Nature, 470(7335), 479–485.

Johnston, R., Jones, K., & Manley, D. (2018). Confounding and collinearity in regression analysis: A cautionary tale and an alternative procedure, illustrated by studies of British voting behaviour. Quality & Quantity, 52, 1957–1976.

Kellermann, V., Overgaard, J., Hoffmann, A. A., Fløjgaard, C., Svenning, J.–C., & Loeschcke, V. (2012). Upper thermal limits of *Drosophila* are linked to species distributions and strongly constrained phylogenetically. Proceedings of the National Academy of Sciences, 109(40), 16228–16233.

Krogh, A. (1916). The respiratory exchange of animals and man (Vol. 13). Longmans, Green.

Lande, R., & Arnold, S. J. (1983). The measurement of selection on correlated characters. Evolution, 1210–1226.

Lighton, J. R. (1998). Notes from underground: Towards ultimate hypotheses of cyclic, discontinuous gas–exchange in tracheate arthropods. American Zoologist, 38(3), 483– 491.

Lighton, J. R. (2018). Measuring metabolic rates: A manual for scientists. Oxford University Press.

Lighton, J. R., & Lovegrove, B. G. (1990). A temperature–induced switch from diffusive to convective ventilation in the honeybee. Journal of Experimental Biology, 154(1), 509–516.

Mallard, F., Nolte, V., Tobler, R., Kapun, M., & Schlötterer, C. (2018). A simple genetic basis of adaptation to a novel thermal environment results in complex metabolic rewiring in Drosophila. Genome Biology, 19(1), 1–15.

Marais, E., Klok, C. J., Terblanche, J. S., & Chown, S. L. (2005). Insect gas exchange patterns: A phylogenetic perspective. Journal of Experimental Biology, 208(23), 4495–4507.

Matthews, P. G., & White, C. R. (2011). Discontinuous gas exchange in insects: Is it all in their heads? The American Naturalist, 177(1), 130–134.

Melna, P. A., & Schwarz, M. P. (1994). Behavioural specialization in pre–reproductive colonies of the allodapine bee *Exoneura bicolor* (Hymenoptera: Anthophoridae). Insectes Sociaux, 41, 1–18.

Messamah, B., Kellermann, V., Malte, H., Loeschcke, V., & Overgaard, J. (2017). Metabolic cold adaptation contributes little to the interspecific variation in metabolic rates of 65 species of Drosophilidae. Journal of Insect Physiology, 98, 309–316.

Monteith, J., & Unsworth, M. (2013). Principles of environmental physics: Plants, animals, and the atmosphere. Academic Press.

Oladipupo, S. O., Wilson, A. E., Hu, X. P., & Appel, A. G. (2022). Why do insects close their spiracles? A meta–analytic evaluation of the adaptive hypothesis of discontinuous gas exchange in insects. Insects, 13(2), 117.

Pettersen, A. K., Hall, M. D., White, C. R., & Marshall, D. J. (2020). Metabolic rate, context-dependent selection, and the competition-colonization trade-off. Evolution Letters, 4(4), 333–344.

Pinheiro, J., Bates, D., DebRoy, S., Sarkar, D., Heisterkamp, S., Van Willigen, B., & Maintainer, R. (2017). Package ‘nlme.’ *Linear and Nonlinear Mixed Effects Models*, Version, 3(1).

R Development Core Team. (2019). R: a language and environment for statistical computing [Computer software].

Roff, D. A., & Fairbairn, D. J. (2012). A test of the hypothesis that correlational selection generates genetic correlations. Evolution: International Journal of Organic Evolution, 66(9), 2953–2960.

Rowe, T. T., Gutbrod, M. S., & Matthews, P. G. (2022). Discontinuous gas exchange in Madagascan hissing cockroaches is not a consequence of hysteresis around a fixed PCO2 threshold. Journal of Experimental Biology.

Schimpf, N. G., Matthews, P. G., & White, C. R. (2012). Cockroaches that exchange respiratory gases discontinuously survive food and water restriction. Evolution: International Journal of Organic Evolution, 66(2), 597–604.

Svensson, E. I., Arnold, S. J., Bürger, R., Csilléry, K., Draghi, J., Henshaw, J. M., Jones, A. G., De Lisle, S., Marques, D. A., & McGuigan, K. (2021). Correlational selection in the age of genomics. Nature Ecology & Evolution, 5(5), 562–573.

Terblanche, J. S., Clusella–Trullas, S., & Chown, S. L. (2010). Phenotypic plasticity of gas exchange pattern and water loss in *Scarabaeus spretus* (Coleoptera: Scarabaeidae): Deconstructing the basis for metabolic rate variation. Journal of Experimental Biology, 213(17), 2940–2949.

Terblanche, J. S., White, C. R., Blackburn, T. M., Marais, E., & Chown, S. L. (2008). Scaling of gas exchange cycle frequency in insects. Biology Letters, 4(1), 127–129.

White, C. R., Alton, L. A., & Frappell, P. B. (2012). Metabolic cold adaptation in fishes occurs at the level of whole animal, mitochondria and enzyme. Proceedings of the Royal Society B: Biological Sciences, 279(1734), 1740–1747.

White, C. R., Blackburn, T. M., Terblanche, J. S., Marais, E., Gibernau, M., & Chown, S. L. (2007a). Evolutionary responses of discontinuous gas exchange in insects. Proceedings of the National Academy of Sciences, 104(20), 8357–8361.

White, C. R., Blackburn, T. M., Terblanche, J. S., Marais, E., Gibernau, M., & Chown, S. L. (2007b). Evolutionary responses of discontinuous gas exchange in insects. Proceedings of the National Academy of Sciences, 104(20), 8357–8361.

Wickham, H. (2011). Ggplot2. Wiley Interdisciplinary Reviews: Computational Statistics, 3(2), 180–185.

Wilke, C. O., Wickham, H., & Wilke, M. C. O. (2019). Package ‘cowplot.’ Streamlined Plot Theme and Plot Annotations for ‘ggplot2.

Winwood–Smith, H. S., & White, C. R. (2018). Short-duration respirometry underestimates metabolic rate for discontinuous breathers. Journal of Experimental Biology, 221(14), jeb175752.

Wohlschlag, D. E. (1960). Metabolism of an Antarctic fish and the phenomenon of cold adaptation. Ecology, 41(2), 287–292.

