## Supplementary material for "Physiological traits and their relationships vary along an elevational gradient within and among Fijian bee species"

### Supplementary Figures

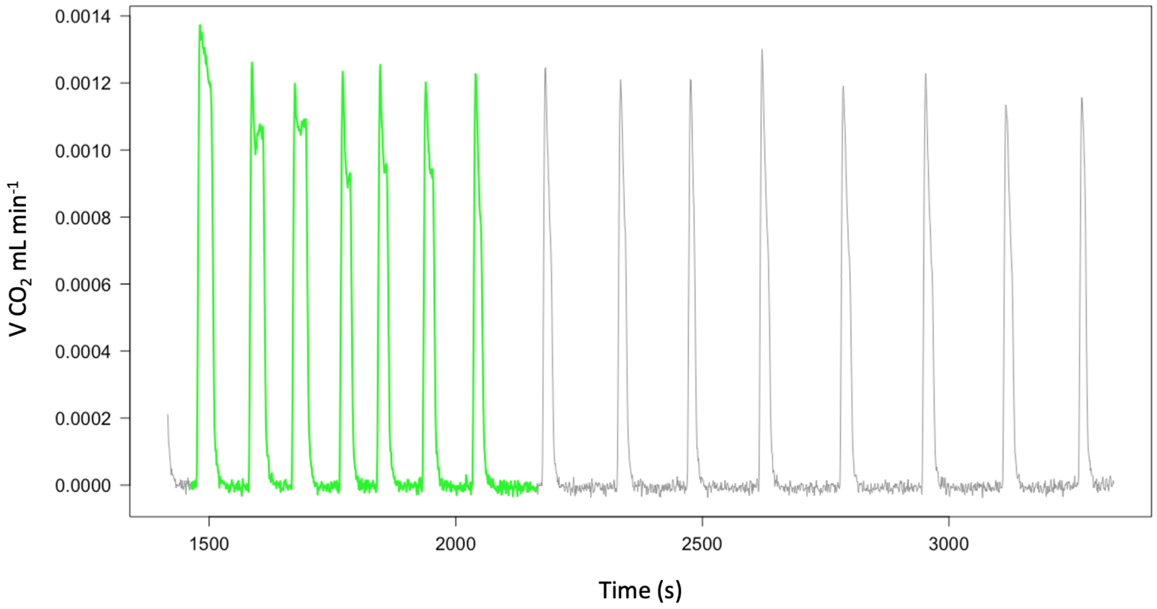


**Supplementary Figure 1.** An example recording of 7 full discontinuous gas exchange cycles ($VCO_2$, ml min-1) (highlighted in green) by an individual bee.

**Box 1. Expansion of hypothesis rationale for model curation.**

Hypotheses to explain variation in **metabolic rate** ($\log_{10}$ transformed to satisfy linear mixed model requirements):

1. Individuals from colder environments will have higher metabolic rates than individuals from warmer environments. MCA supported.
2. Individuals from low precipitation environments have the lowest metabolic rates to avoid respiratory water loss. MCA rejected.
3. Drying power of the air (VPD) best explains variation in metabolic rate, where individuals from high VPD regions (warm and dry) have the lowest metabolic rates to decrease respiratory water loss. MCA rejected.
4. Due to the 'wet' and 'dry' side of Viti Levu (Figure 1c), temperature and precipitation independently impact metabolic rate, where both low temperatures and high precipitation select for low metabolic rates. However, thermal environments can vary in their level of precipitation depending on location (wet or dry side and altitude). MCA-like pattern is observed, but precipitation is also an important predictor of metabolic rate.
5. Null hypothesis - Metabolic rate changes across altitude, but some other abiotic variable is shaping variation in metabolic rate. MCA rejected.

Hypotheses to explain variation in **frequency of gas exchange**:

6. Individuals from low precipitation environments will have the lowest frequencies of gas exchange to reduce water loss. Hygric hypothesis supported.
7. Environmental temperature impacts frequency of gas exchange due to the direct thermodynamic effects of temperature on performance (Angilletta Jr et al., 2010), where rates of gas exchange increase with warming temperatures. Hygric hypothesis rejected.
8. Drying power of the air (VPD) best explains variation in frequency of gas exchange, where individuals from high VPD environments have the lowest frequency of gas exchange to decrease respiratory water loss. Hygric hypothesis supported.
9. Temperature and precipitation independently shape variation in frequency of gas exchange, where both high temperatures and low precipitation select for low frequencies of gas exchange, but locations can vary in their temperature/precipitation combination. Hygric hypothesis supported, but temperature also plays an important role in explaining variation in frequency of gas exchange.
10. Null hypothesis - frequency of gas exchange changes across altitude, but some other abiotic variable (other than temperature, precipitation and aridity) is shaping variation in frequency of gas exchange.

Hypothesis to explain **co-variation between metabolic rate and frequency of gas exchange**:

11. The relationship between metabolic rate and frequency of gas exchange will be impacted by the environmental temperature species inhabit, where the frequency of gas exchange - metabolic rate relationship will be steeper in cooler

environments than warmer environments as selective pressure to increase metabolic rate in cold environments will be greater than the selective pressure to decrease frequency of gas exchange.

12. The relationship between metabolic rate and frequency of gas exchange will be impacted by aridity (VPD), where the ventilation rate - metabolic rate relationship will be steeper in low VPD environments as selective pressure on ventilation rate will be relaxed in low VPD environments.
13. The relationship between metabolic rate and ventilation rate will be impacted by precipitation, where selective pressure to reduce ventilation rate will be greater in dry environments than the selective pressure to reduce metabolic rate.
14. The relationship between metabolic rate is best explained by the additive effects of temperature and precipitation, where both warm temperatures and low precipitation independently select for low metabolic rates and ventilation rates.
15. Null hypothesis - an interaction between metabolic rate and altitude better explains variation in ventilation rate than the other models, suggesting that another variable better explains the covariation between metabolic rate and ventilation rate.

---

### Supplementary Tables

**Supplementary Table 1.** Summary of the abiotic variable ranges that each species was collected across.

| **Abiotic variables** | ***Braunsapis puangensis*** | ***Homalictus fijiensis*** | ***Homalictus groomi*** | ***Homalictus tuiwawae*** |
|---|---|---|---|---|
| Minimum altitude | 6.0 | 6.0 | 858.0 | 575.0 |
| Maximum altitude | 693.0 | 1072.0 | 922.0 | 922.0 |
| Altitudinal range | 687.0 | 1066.0 | 64.0 | 347.0 |
| Mean temp. July minimum | 20.9 | 18.6 | 18.6 | 19.9 |
| Mean temp. July maximum | 22.9 | 23.1 | 19.9 | 20.4 |
| July temp range | 2.1 | 4.5 | 1.3 | 0.5 |
| Mean precipitation minimum | 87.1 | 57.6 | 103.0 | 103.0 |
| Mean precipitation maximum | 132.2 | 187.8 | 187.8 | 105.7 |
| Precipitation range | 45.1 | 130.1 | 84.8 | 2.8 |
| Mean VPD July minimum | 0.32 | 0.10 | 0.10 | 0.22 |
| Mean VPD July maximum | 0.56 | 0.56 | 0.22 | 0.27 |
| VPD range | 0.24 | 0.46 | 0.12 | 0.05 |
| **Sample size** | **44** | **125** | **16** | **23** |

**Supplementary Table 2.** Testing the MCA hypothesis within *Homalictus fijiensis*. Best fitting model summary (lowest AIC from strong inference model comparison) showing the effect of body mass, average temperature of the coldest month and average precipitation of the driest month on routine metabolic rate.

| Coefficient | Estimate | SE | df | t-value | P-value |
|---|---|---|---|---|---|
| Intercept | -3.09 | 0.30 | 92 | -10.59 | < 0.001 |
| $Log_{10}$(body mass) | 0.76 | 0.07 | 92 | 11.57 | < 0.001 |
| **$T_{min}$** | **-0.041** | **0.04** | **92** | **-3.29** | 0.0014 |
| **$P_{dry}$** | **-0.0017** | **0.002** | **92** | **-3.63** | < 0.001 |

**Supplementary Table 3.** Testing the MCA hypothesis across species. Best fitting model summary (lowest AIC from strong inference model comparison) showing the effect of body mass, species, an interaction between sex and species, average temperature of the coldest month and average precipitation of the driest month on routine metabolic rate.

| Coefficient | Estimate | SE | df | t-value | P-value |
|---|---|---|---|---|---|
| Intercept | -3.16 | 0.33 | 167 | -9.65 | < 0.001 |
| log10(mass) | 0.76 | 0.07 | 167 | 11.23 | < 0.001 |
| *Homalictus fijiensis* | -0.007 | 0.04 | 167 | -0.17 | 0.86 |
| *Homalictus groomi* | -0.19 | 0.07 | 167 | -2.63 | 0.009 |
| *Homalictus tuiwawae* | -0.002 | 0.07 | 167 | -0.03 | 0.978 |
| **$T_{min}$** | **-0.037** | **0.013** | 167 | **-2.90** | **0.004** |
| **$P_{dry}$** | **-0.001** | **0.0004** | 167 | **-3.45** | **0.0007** |

**Supplementary Table 4.** Testing the hygric hypothesis within *Homalictus fijiensis*. Best fitting model summary (lowest AIC from strong inference model comparison) showing the effect of body mass, sex, altitude, and average precipitation of the driest month on ventilation rate.

| Coefficient | Estimate | SE | df | t-value | P-value |
|---|---|---|---|---|---|
| Intercept | 2.55 | 0.35 | 92 | 7.14 | <0.001 |
| $Log_{10}$(body mass) | -0.51 | 0.09 | 92 | -5.76 | <0.001 |
| $Log_{10}$(MR) | 0.67 | 0.09 | 92 | 7.74 | <0.001 |
| Altitude | -0.00008 | 0.00004 | 92 | -1.72 | 0.089 |

**Supplementary Table 5.** Testing the hygric hypothesis among bee species. Best fitting model summary (lowest AIC from strong inference model comparison) showing the effect of body mass, sex, altitude, and average precipitation of the driest month on ventilation rate.

| Coefficients | Estimate | SE | DF | t-value | P-value |
|---|---|---|---|---|---|
| Intercept | 1.99 | 0.26 | 167 | 7.78 | <0.001 |
| log10(mass) | -0.43 | 0.07 | 167 | -5.96 | <0.001 |
| Homalictus fijiensis | 0.066 | 0.03 | 167 | 1.92 | 0.056 |
| *Homalictus groomi* | -0.029 | 0.06 | 167 | 0.45 | 0.65 |
| *Homalictus tuiwawae* | 0.095 | 0.06 | 167 | 1.55 | 0.12 |
| $Log_{10}$MR | 0.55 | 0.06 | 167 | 9.05 | <0.001 |
| Altitude | -0.00005 | 0.00004 | 167 | -1.16 | 0.248 |

**Supplementary Table 6**. Assessing how frequency of gas exchange and metabolic rate co-vary across abiotic gradients across bee species. Best fitting model summary (lowest AIC from strong inference model comparison) coefficients.

| Coefficients | Value | SE | df | t-value | P-value |
|---|---|---|---|---|---|
| Intercept | 1.01 | 0.07 | 2 | 14.59 | 0.005 |
| Altitude | -0.0008534 | 0.0001 | 2 | -7.90 | 0.016 |
